# Taxonomy Analysis in Bacterial Kingdom based on Protein Domain: A Comparison Study

**DOI:** 10.1101/2021.09.17.460715

**Authors:** Boqian Wang, Jianglin Zhou, Yuan Jin, Mingda Hu, Yunxiang Zhao, Xin Wang, Haoyi Yang, Xingfei Gong, Fengwei Zhang, Zehan Zhang, Fuqiang Kang, Long Liang, Junjie Yue, Hongguang Ren

**Author notes:** Correspondence: Hongguang Ren or Junjie Yue or Long Liang. These authors contributed equally to this paper (Boqian Wang, Jianglin Zhou). **Author Contributions:** HR, LL, and JY formulated the study; BW and JZ performed the research and analyzed the data; YJ, MH, YZ, XW, HY, XG, FZ, ZZ, and FK participated in analysis and discussion; BW and HR drafted the manuscript; All authors read and approved the final manuscript.

## Abstract

The taxonomy research on the bacterial kingdom is important and can be usually conducted by the analysis on conserved genes, 16s rRNA, protein domain, and so on. Comparatively, the protein domains maintain a most direct relationship with phenotypes. In this paper, based on the protein domain, we propose a 3-step framework to standardize the classification process of bacteria. Different model candidates are involved and discussed in each step. By comparing the classification results with existing taxonomy, we select the most appropriate candidate to improve the framework, and furthermore, discuss their biological significance. Finally, we put forward taxonomy suggestions based on the best classification results.

**Significance Statement:** We standardize a 3-step framework to carry out bacterial taxonomy research based on protein domain. Furthermore, we filter out the best solution in each step that can together generate the most appropriate classification result, and at the same time, we discuss the biological significant it indicates. Finally, we propose suggestions on NCBI bacterial taxonomy based on the classification results.

## Introduction

Traditional classification method is based on the phenotypes, indicating that the previous taxonomy results mainly reflect the commons of phenotypes. With the development of biotechnology, the genetic information offers us the chance to look into the essence of taxonomy.

The comparison of genome sequences is the most intuitive method which directly focuses on the original genome data [1, 2]. Comparatively, the protein sequence can mask some gene-level differences for the reason that the same amino acid could be translated from different codons [3, 4]. Furthermore, the protein domain, as the basic functional unit of protein, is a bridge that can directly connect genetic information with phenotypes [5-7]. It is inferred from a given sequence of amino acids, the length of which is usually between 50 to 350.

These methods can be separately applied to scenarios with different requirements. For example, to investigate the gene mutation or recombination events in coronavirus, genome level sequences should be compared and analyzed [8, 9]. Whereas when it comes to a larger scope, like a kingdom or a phylum, the species usually have relatively far relationship. In this case, it is difficult to use the whole genome to infer phylogeny due to the complex evolutionary background of the species, like the recombination or horizontal gene transfer events. Besides, utilizing conserved house-keeping genes solo for classification may lost the phenotype information of species. Comparatively, protein domain-based method can serve as a compromise for both phylogeny and phenotypes, which is perfect for taxonomy analysis of bacteria.

Nowadays, many taxonomy researches are conducted based on the protein domain [5-7]. However, several important problems in this area still remain to be solved. First, many domain-based methods have been individually proposed and applied without systematical analysis and comparison. Secondly, the biologic significance and the rationality of the methods are usually missed.

In this paper, we propose a protein domain-based framework to classify species examples from bacterial kingdom. The framework consists of three steps, which separately involve statistic model of domains, distance model, and construction and analysis models of taxonomy tree. We design different model candidates in each step for comparison. On the one hand, we try to find the best model solution in each step as well as the biological significance it indicates. It helps us understand the relationship between protein domains and the phenotypes. On the other hand, based on the classification result, we propose some taxonomy suggestions, most of which can be further verified by the related works.

## Materials and Methods

The source codes (in Python) related to this paper can be found on GitHub: https://github.com/wr-sky/Domain-Bac-Tax.

### Genome selection

Genome sequences of the whole bacterial domain are downloaded according to the metadata (Feb. 7. 2021) on NCBI website: ftp://ftp.ncbi.nlm.nih.gov/genomes/README_assembly_summary.txt.

Totally, 205791 species are recorded in the text file. For a more credible result, we select the sequences that are “complete genome” in “assembly_level” column, “latest” in “version_status” column, “full” in “genome_rep” column, and “representative genome” or “reference genome” in “refseq_category” column. Finally, we downloaded 2587 sequences in faa (FASTA Amino Acid) and fna (FASTA Nucleic Acid) formats from NCBI. The quality of each sequence (fna format) is then inspected by the CheckM program with the standard shown in Equation (1).

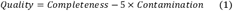

Two sequences with quality results under 95% are removed, and thus finally 2568 genome sequences are utilized for further analysis. Their information can be found in Supplementary Information (Dataset S1). Besides, 6 species from Archaea domain are involved as the external species, which are: Desulfurococcus amylolyticus, Halorhabdus utahensis, Halomicrobium mukohataei, Halogeometricum borinquense, Nitrososphaera viennensis, and Saccharolobus solfataricus.

### Protein domain dataset

The 2568 nucleic acid sequences in FASTA format are analyzed by the pfam_scan.pl program with default sets [10]. The results in tsv-format output file list the possibly existed domains in each sequence. Domains with overlapped region will be polished by selecting the one with the maximal bit score value. Then the protein domain dataset will be handled by our proposed framework below.

### Structure of framework

As shown in Figure 1, our proposed framework consists of three steps, which involve the statistic model of domains, the distance model, and the construction and analysis models of taxonomy tree. The statistic model of domains collects the domain information from the protein domain dataset. The distance model is responsible to define the distance of each pair of species according to the statistic results of domains. Finally, the taxonomy tree is constructed and analyzed based on the distance results.

**Figure 1.**
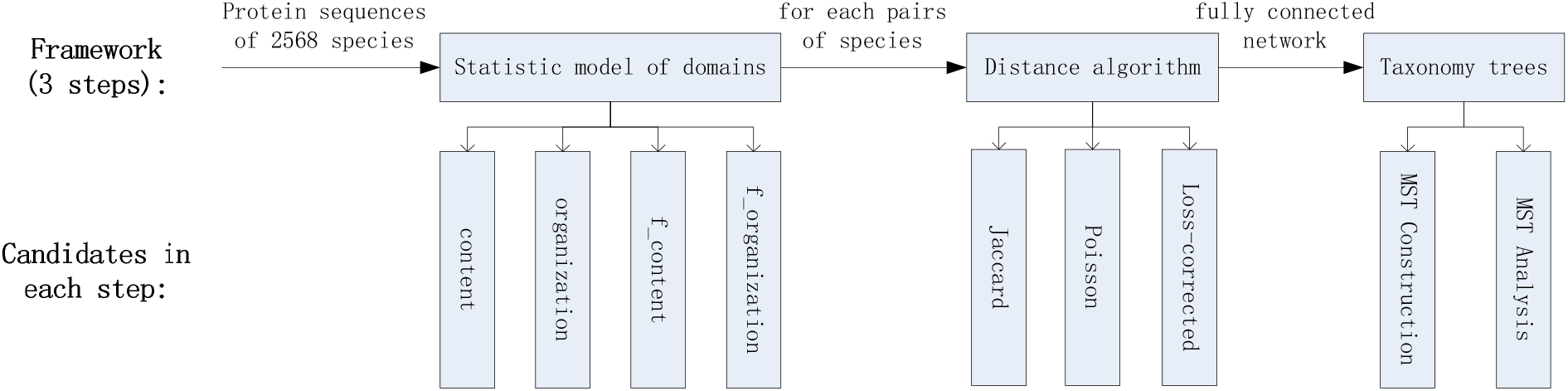
3-step process in our proposed framework and candidates in each step.

#### Statistic models of domains

In the first step, we propose four statistic models of domains. As shown in Figure 2, in the first statistic model, the “content” model records the domain content in the species. It emphasizes the importance of individual function of each domain and only considers the presence or absence of a domain. In the second model, the “organization” model takes the sequence of domains in a protein into consideration. Compared with the “content” model, it focuses on the sequence and thus, the co-function of all domains in a protein.

**Figure 2.**
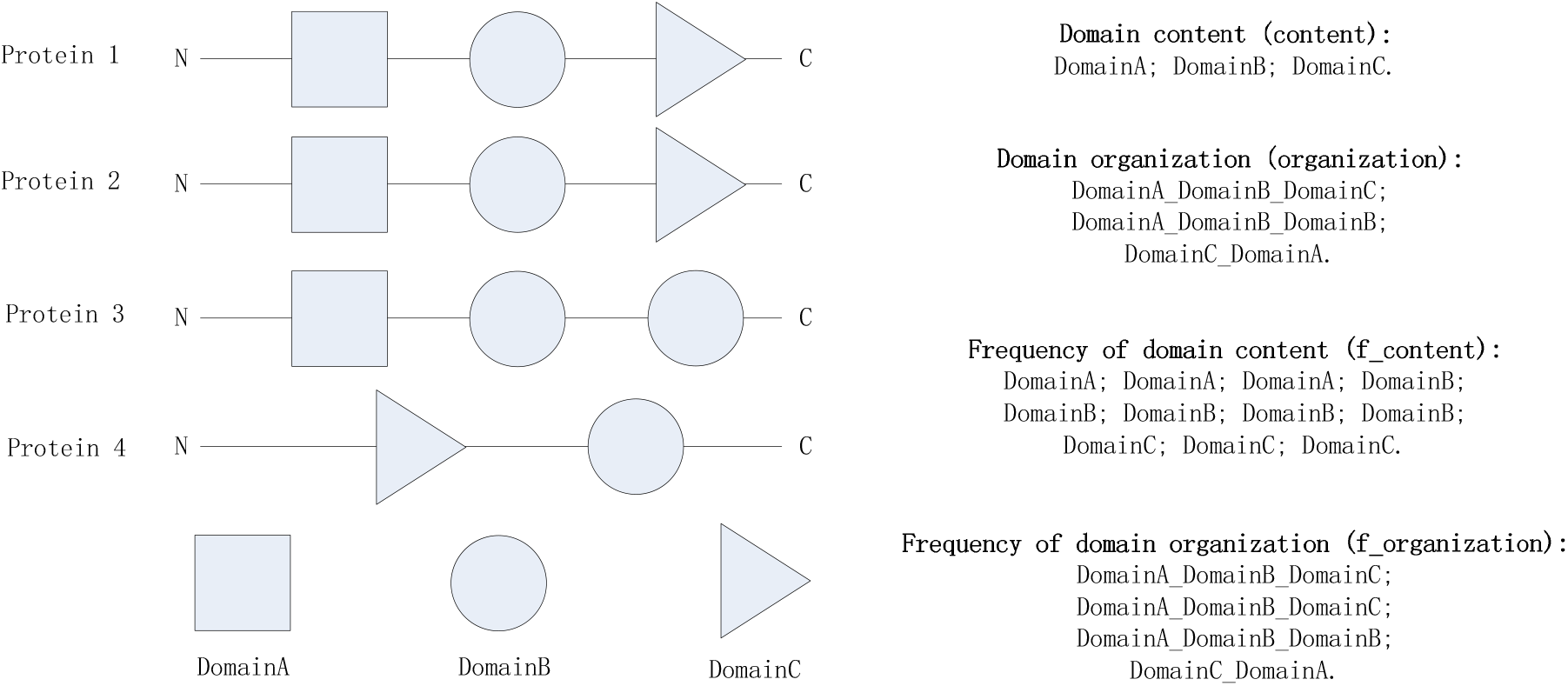
An example of domains in one species and the corresponding records separately by four different statistic models.

Besides, by additionally considering the frequency of a domain in the species, we design another two statistic models, namely the “f_content” model and the “f_organization” model, which are separately corresponding to the “content” model and the “organization” model.

- By comparing the “content” and “organization” models, we try to find out whether the *single domain* or the *domains organization* can influence the taxonomy and the bacterial phenotypes to the greatest extent.
- The adoption of frequency aims to explore the influence of the number of domains (“content” or “organization”) on bacteria. That is to answer the question: Which is more important when deciding bacterial phenotypes in the evolutionary history, the presence of a domain or the number of a domain?

To reduce the influence of redundant information on the final taxonomy result, other models such as the one that combines domain content and domains organization together will be not considered as a model candidate.

Finally, four statistic models will be compared in the *Framework Results* section to find out the best solution as well as the answers to the two questions above.

#### Distance models

In the second step, we involve the distance models that calculate the distance of each pair of species based on domain statistic results. Three methods are compared and discussed, which are commonly utilized in this area and can reflect the biological significance from different perspectives [5][7].

The first distance model is the Jaccard distance as shown in Equation (2). Parameters a, b, and c are separately the number of domains in species A, species B, and commonly in species A and B as shown in Figure 3. The concept of domain here could represent domain content or domains organization.

**Figure 3.**
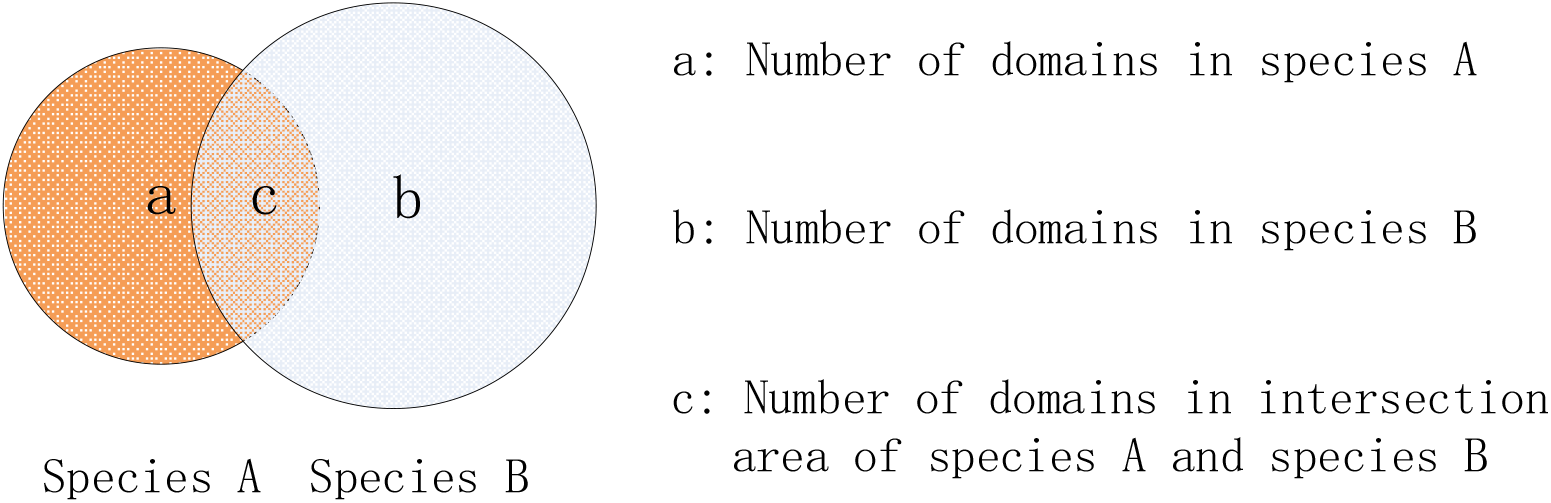
A Venn diagram that shows concept of the corresponding three parts when comparing the domains of two species.

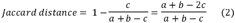

Jaccard distance is a very common way to calculate the similarity/differences of two sets. When it comes to our scenario, Jaccard distance deduces the evolutionary distance of two species under the assumption that the change of domain (by mutation, loss or recombination) happens randomly and independently.

The second distance model is the Poisson distance as shown in Equation (3). Different from Jaccard distance, it is under the assumption that the change of domain follows the Poisson process [7]. 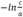 and 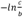 represent the evolutionary distances between the two species and their ancestor. The evolutionary distance between the two species is defined as the geometric mean of their distances to the common ancestor.

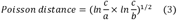

The third distance model is the Loss-corrected distance as shown in Equation (4). It considers the possibility of massive gene loss during evolutionary history of bacteria. Thus, to reduce its influence, the distance is corrected by utilizing the smaller domain set as the standard.

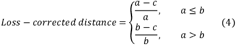

The values calculated by the three models range from 0 to 1 with 1 implying the farthest evolutionary distance and 0 the nearest. This will result in a fully connected network with each node representing a species and each line representing the evolutionary distance.

These three distance models will also be compared in the *Framework Results* section to find out the best solution as well as the fact that which assumption can fit the real evolutionary process to the greatest extent.

4 statistic models together with 3 distance models can finally form 12 different combinations, representing 12 different processing methods in the framework.

#### Construction and analysis models of taxonomy tree

In the third step, the fully connected network will be polished to Minimum-cost Spanning Tree (MST) by Prim’s algorithm as shown in Figure 4. First, it will randomly select one species as the initial node of the MST. Then, the MST involves another node which has the minimal distance with a node already in the MST and connects them together. The second step repeats until all nodes are involved in the MST. According to the Prime’s algorithm, the MST try to find the pair of nodes with the minimal distance in each step. Reflected to our scenario, the MST will connect the species with shortest evolutionary distance together.

**Figure 4.**
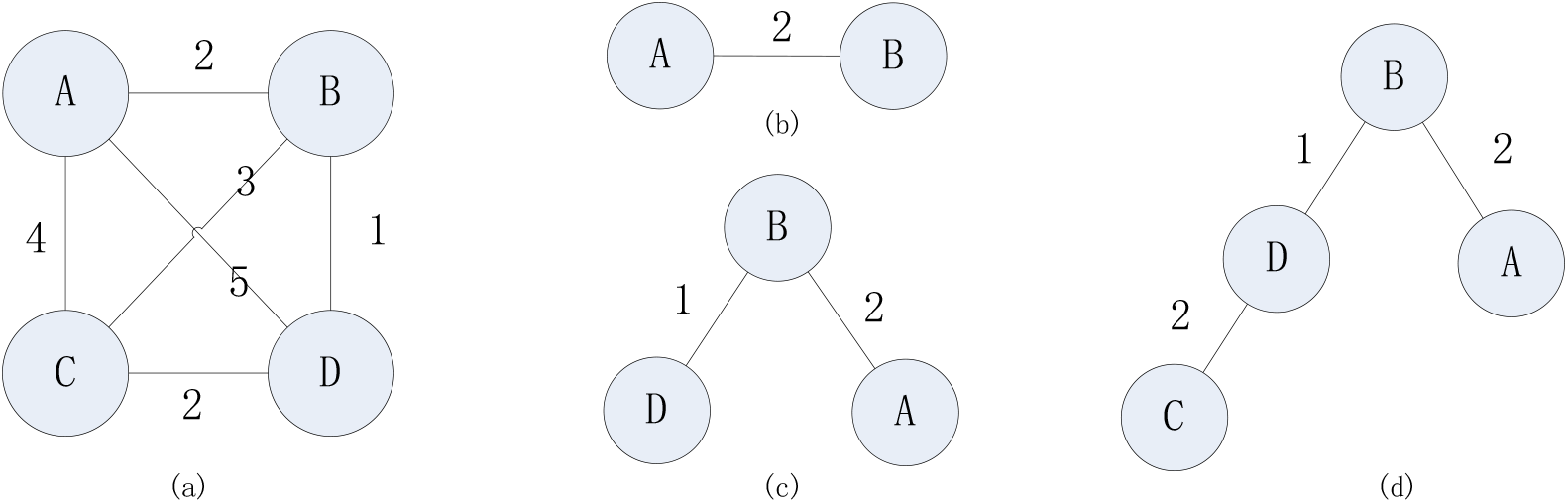
An example to construct the MST based on the fully connected network. (a) A fully connected network with distances marked. (b) Node A is initially selected and then its nearest node B is connected to Node A. (c) Node D is connected to node B since it has the nearest distance to node A or node B compared with node C.(d) Node C is nearer to D compared with the distance to node B or node A, and thus node C is finally connected to node D.

We write a program to cluster and compare the MST results according to the NCBI taxonomy database, which mainly focuses on three aspects below:1. The number of species in each phylum that are separated from their main part.

2. The number of isolated parts for each phylum.

3. The separated species that totally exist in the MST.

The comparison results in the *Framework Results* section can confirm the feasibility and effectiveness of the adoption of MST in our framework.

## Framework Results

12 MST results generated by 12 different methods in the framework can be found in Supplementary Information (Figure S1 ∼ Figure S12). Taking the “content” model with Jaccard distance model as an example, we show the MST result in Figure 5. For ease of analysis, the figure is manually colored in phylum and class levels according to the NCBI taxonomy database. It clearly shows that the result matches the NCBI taxonomy very well, especially in the phylum level.

**Figure 5.**
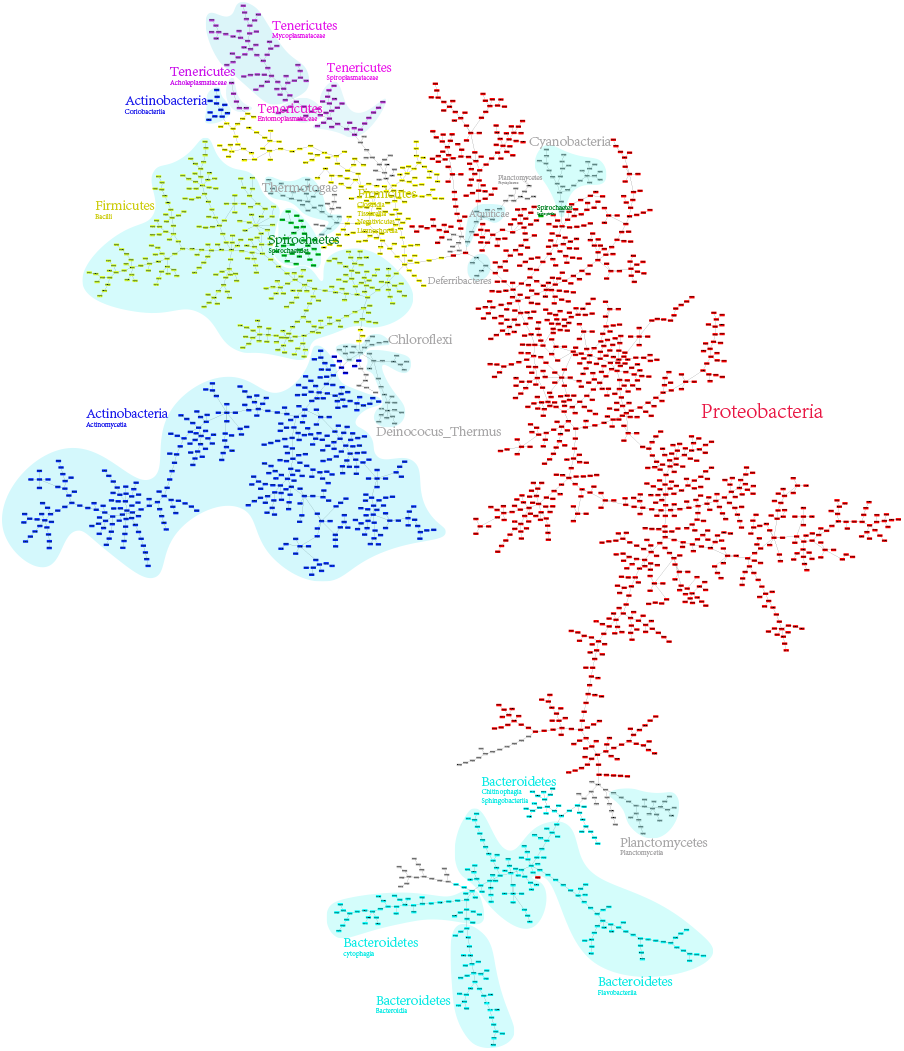
MST result by “content” model and Jaccard distance model. The MST result is manually preprocessed by grouping species in the phylum and class levels, and the phyla including more than 10 species are distinguished with different colors.

We write a program to quantify the results, the details of which can be found in Supplementary Information (Table S1 & Table S2). Table S1 shows the results of each phylum with Jaccard and Poisson distance models, which list the number of species in each small group that has been isolated from its main part (the group with the maximal number of species). If no species is isolated for a phylum, the corresponding record will remain blank.

Since the Loss-corrected distance model results in too many isolated small groups for some phyla, Table S2 only records the number of groups that each phylum has been isolated into with the Loss-corrected distance model.

To compare the results more clearly, we propose three standards: the percentage of isolated species (arithmetic percentage), the weighted percentage of isolated species (weighted percentage), and the number of phyla possessing more than one group (phyla number). As shown in Equation (7) and (8), the parameter *S*_*i*_ stands for the number of isolated species in each phylum and *T*_*i*_ stands for the number of species in each phylum. The parameter *i* ranges from 1 to 31, representing 30 bacterial phyla and 1 archaea domain.

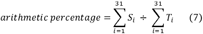

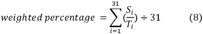

We show the results in Figure 6 and mark out the best and the second-best solutions for each standard with red frames. It is obvious that org_ja (“organization” mode & Jaccard distance model) and org_po (“organization” mode & Poisson distance model) can always produce better results under each standard, and furthermore, org_ja is slightly better than org_po.

**Figure 6.**
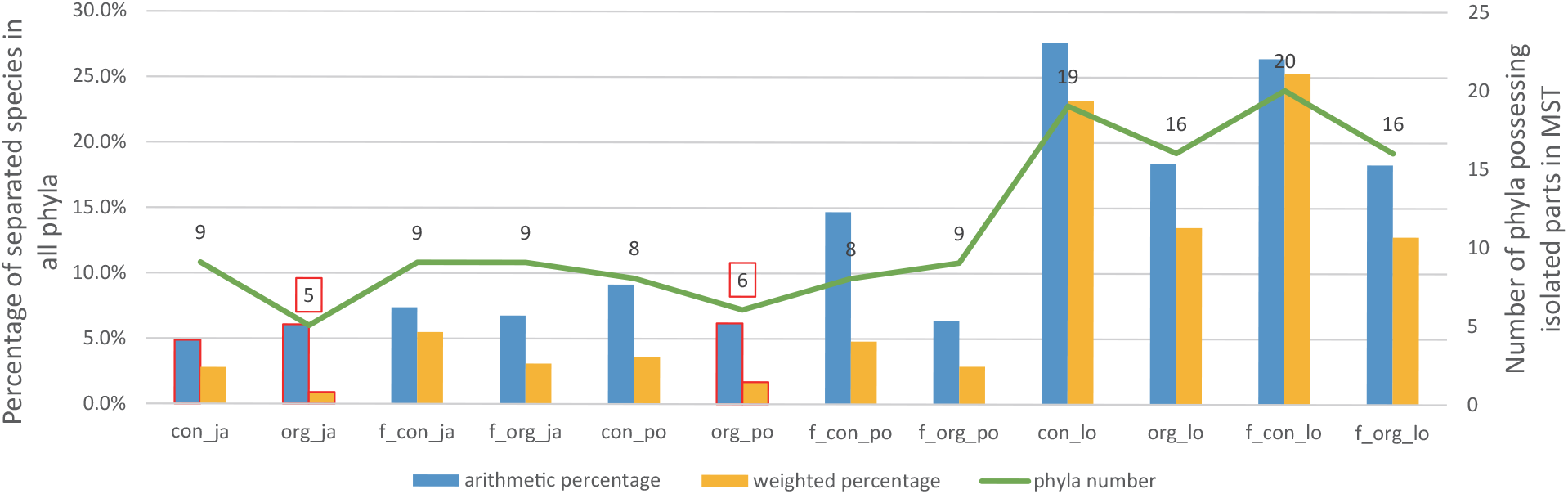
Twelve methods are compared by three different standards. Con, org, f_con, and f_org separately represent “content” model, “organization” model, “f_content” model, and “f_organization” model. Ja, po, and lo stand for Jaccard, Poisson, and Loss-corrected distance models, respectively.

The results show that the “organization” model is better than the other three statistic models in most cases. On the one hand, it indicates that the *domains organization* plays a more important role to influence the taxonomy and the bacterial phenotypes compared with the *single domain*.

On the other hand, we can get the conclusion that the number of a domain organization in a species is less important that the presence of a domain organization.

As for the distance models, Jaccard distance is slightly better than the Poisson distance in most cases, both of which are significantly better than the Loss-corrected distance. This may be due to the reason that the evolutionary distance calculated by Loss-corrected algorithm is decided by the species with smaller genome size to a large extent. It can also explain the phenomenon that many species are easily to form the center of a circle in MST under Loss-corrected distance model. To this end, we can get the conclusion that the evolution process of domain in bacteria is usually random or follow Poisson process without high-frequent massive gene loss.

## Taxonomy Discussion

To compare the classification results with NCBI taxonomy database, we abstract the MST generated by the org_ja process into Figure 7. It has been clearly shown that five groups of species are far away separated from their main phylum parts, which, marked by the yellow background in Figure 7, are Actinobacteria (Coribobacteriia), Tenericutes (Acholeplasmatales), Spirochaetes (Leptospirales), Planctomycetes (Phycisphaerae), and Proteobacteria (Glaciecola amylolytica), respectively. It indicates a relatively high protein domain differences with the other species in the same phylum. Besides, the Proteobacteria phylum is isolated into four parts by three phyla (Aquificae, Thermodesulfobacteria, and Deferribactetes) that only contain very small number of species examples. Thus, we will mainly look into details of the corresponding 19 species that are far away isolated from their main parts. Their taxonomy information is listed in Table 1.

**Table 1.** The nineteen species include eleven species in Actinobacteria, five species in Tenericutes, and one species in Planctomycetes, Proteobacteria and Spirochaetes respectively. The eleven Actinobacteria species are separated from the main Actinobacteria species in ‘class’ level, which means that if and only if the species in the class Coriobacteriia are separated from the species in the other classes of the phylum Actinobacteria. The separation level of each phylum is marked out in green words.

| GCF | Phylum | Class | Order | Family | Genus | Species |
| --- | --- | --- | --- | --- | --- | --- |
| 900169485 | Actinobacteria | Coriobacteriia | Eggerthellales | Eggerthellaceae | Phoenicibacter | Phoenicibacter |
| 000023845 | Actinobacteria | Coriobacteriia | Eggerthellales | Eggerthellaceae | Cryptobacterium | Cryptobacterium |
| 900637575 | Actinobacteria | Coriobacteriia | Eggerthellales | Eggerthellaceae | Slackia | Slackia |
| 001643775 | Actinobacteria | Coriobacteriia | Eggerthellales | Eggerthellaceae | Denitrobacterium | Denitrobacterium detoxificans |
| 000024265 | Actinobacteria | Coriobacteriia | Eggerthellales | Eggerthellaceae | Eggerthella | Eggerthella lenta |
| 000478885 | Actinobacteria | Coriobacteriia | Eggerthellales | Eggerthellaceae | Adlercreutzia | Adlercreutzia equolifaciens |
| 000195315 | Actinobacteria | Coriobacteriia | Coriobacteriales | Coriobacteriaceae | Coriobacterium | Coriobacterium |
| 900119915 | Actinobacteria | Coriobacteriia | Coriobacteriales | Atopobiaceae | Olsenella | Olsenella |
| 000143845 | Actinobacteria | Coriobacteriia | Coriobacteriales | Atopobiaceae | Olsenella | Olsenella uli |
| 000024225 | Actinobacteria | Coriobacteriia | Coriobacteriales | Atopobiaceae | Lancefieldella | Lancefieldella |
| 003966955 | Actinobacteria | Coriobacteriia | Coriobacteriales | Atopobiaceae | Parolsenella | Parolsenella |
| 000967915 | Tenericutes | Mollicutes | Acholeplasmatales | Acholeplasmataceae | Acholeplasma | Acholeplasma |
| 900660755 | Tenericutes | Mollicutes | Acholeplasmatales | Acholeplasmataceae | Acholeplasma | Acholeplasma |
| 000018785 | Tenericutes | Mollicutes | Acholeplasmatales | Acholeplasmataceae | Acholeplasma | Acholeplasma |
| 000968055 | Tenericutes | Mollicutes | Acholeplasmatales | Acholeplasmataceae | Acholeplasma | Acholeplasma |
| 900660745 | Tenericutes | Mollicutes | Acholeplasmatales | Acholeplasmataceae | Acholeplasma | Acholeplasma |
| 000266885 | Spirochaetes | Spirochaetia | Leptospirales | Leptospiraceae | Turneriella | Turneriella parva |
| 003856375 | Proteobacteria | Gammaprot | Alteromonadales | Alteromonadaceae | Glaciecola | Glaciecola |
| 001999965 | Planctomycetes | Phycisphaerae | Sedimentisphaerales | Sedimentisphaeraceae | Limihaloglobus | Limihaloglobus sulfuriphilus |

**Figure 7.**
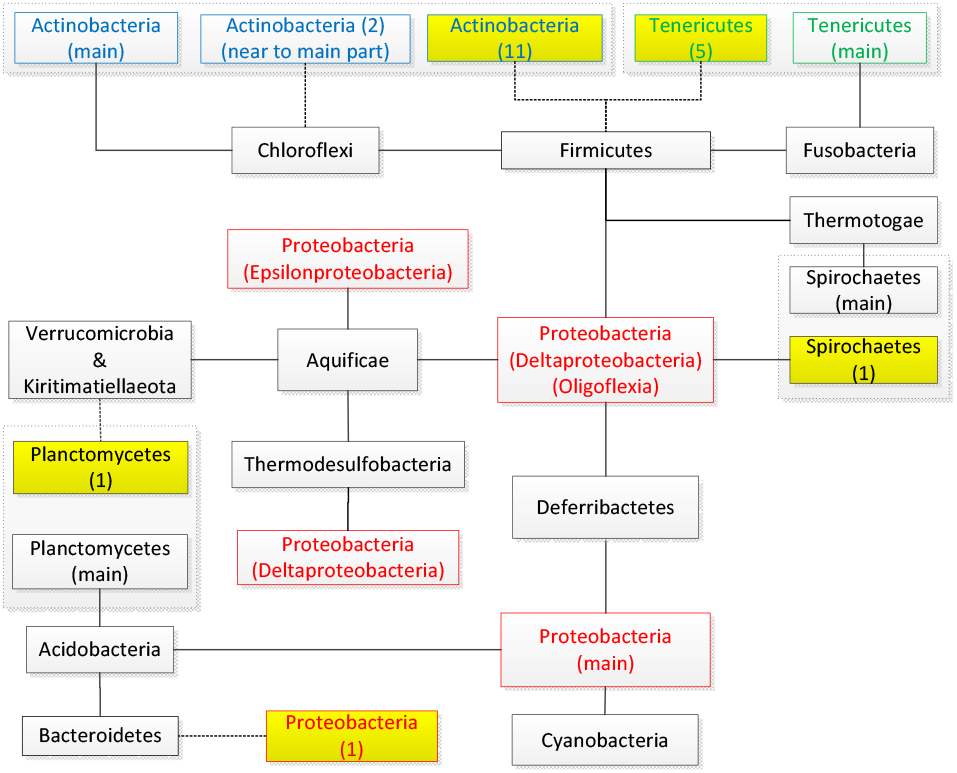
Abstracted figure of MST generated by org_ja method. Six groups of species separated from their main phylum parts are connected to their neighboring phyla by the dashed line. Five of them are far away from their main phylum parts with yellow background and the numbers of species are also marked out.

### Actinobacteria

432 examples of Actinobacteria are included in the dataset and 11 species of them are separated from the main part in the class level, being connected to Firmicutes (Erysipeiotrichia). All these 11 species belong to the class Coriobacteriia. So, we suggest to move the class Coriobacteriia to the phylum Firmicutes. This can be supported by two evidences. On the one hand, in [14], the identification of a number of conserved signature indels (CSIs) and conserved signature proteins (CSPs) shows that they are commonly and uniquely shared by the most members of all other classes of the phylum Actinobacteria, except Coriobacteriia, which branches more deeply than all other actinobacteria. It suggested that the class Coriobacteriia should be excluded from the phylum Actinobacteria. On the other hand, the species Symbiobacterium thermophilum has been moved from the phylum Actinobacteria to Firmicutes by CSIs and CSPs standards, as well as the genome sequence and other lines of evidences [15, 16].

### Tenericutes

94 examples are included in the dataset and 5 species of them are separated from the main phylum part, being connected to Firmicutes (Erysipeiotrichia). The 5 species belong to the same genus and are separated from the other species of the phylum Actinobacteria in the order (Acholeplasmatales) level. Based on our classification results, we suggest to move the order Acholeplasmatales of the phylum Tenericutes to the phylum Firmicutes. There are two evidences to support our suggestion. First, the phylum Tenericutes only contains one single class Mollicutes, which had belonged to Firmicutes [17]. Secondly, the class Mollicutes consists of three orders which are Acholeplasmatales, Entomoplasmatales, and Mycoplasmatales. Due to the reason that the order Acholeplasmatales does not require sterol for growth, which is different from the other species in the phylum Tenericutes, the order Acholeplasmatales maintains a far relationship with the other orders [18].

### Spirochaetes

28 examples are included in the dataset and 1 species of them are separated from the main phylum part, being connected to Proteobacteria (Oligoflexia). This species (order Leptospirales) is separated from the other species (order Spirochaetales) in the phylum Thermotogae in the order level. To erase the deviation caused by the single species example, we additionally involved another 8 species in the order Leptospirales. Their detailed information is listed in Supplementary Information (Dataset S2). The updated MST is shown in Figure 8 (left), where we can find that the order Leptospirales is still isolated from the order Spirochaetales.

**Figure 8.**
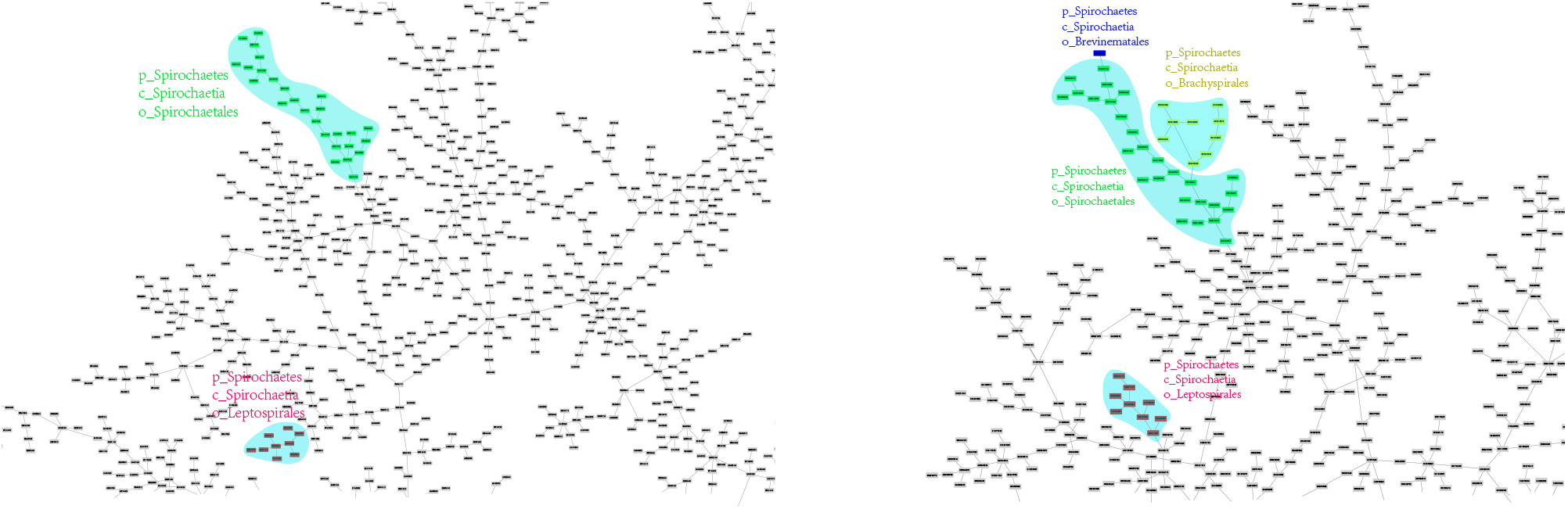
Eight species in the order Leptospirales are added to generate the MST (left). Another nine species in Brachyspirales and one species in Brevinematales are involved to explore the relationship of the four orders in the phylum Spirochaetes (right).

Actually, the order Leptospirales had been a family under the order Spirochaetales by the 16s rRNA analysis results, and has been latterly promoted into an order [17] [19]. The other two orders are Brachyspirales and Brevinematales, respectively. We compare the phenotypes of these four orders as well as the order Oligoflexia (neighbor order in the MST result) in Table 2.

**Table 2.** Four characteristics of five orders are compared. Four orders in the phylum Spirochaetes share the same helical cell shape. The order Leptospirales has no hooked ends which is totally different from the other three orders in the phylum Spirochaetes. Besides, the Periplasmic flagella of Spirochaetales, Brachyspirales and Brevinematales is overlapped and located in the central region of the cell. Finally, Species in Leptospirales are aerobic or at least microaerophilic, which is not the fact in the other three orders in the phylum Spirochaetes, where species are usually anaerobic or microaerophilic at most. As for the order Oligoflexia in the phylum Proteobacteria, it is also quite different from the order Leptospirales among these characteristics.

| Order | Shape | Hooked ends | Periplasmic flagella | Habit |
| --- | --- | --- | --- | --- |
| Spirochaetales | Helical, coccoid | No | Central, overlapped | Anaerobic, microaerophilic |
| Brachyspirales | Helical | No | Central, overlapped | Anaerobic |
| Brevinematales | Helical | No | Central, overlapped | Microaerophilic |
| Leptospirales | Helical | Yes | Central, not overlapped | Aerobic, microaerophilic |
| Oligoflexia[20] | Pleiomorphic | - | No | Aerobic |

Table 2 shows the fact that the order Leptospirales has different characteristics from either the other order in phylum Spirochaetes or its MST-connected order Oligoflexia. To investigate into their relationship, we additionally involve another 1 species in order Brevinematales and 9 species in the order Brachyspirales to create a new MST with org_ja method. The result is shown in Figure 8 (right). The detailed information is listed in Supplementary Information (Dataset S2). The orders Spirochaetales, Brachyspirales and Brevinematales are connected to each other, while the order Leptospirales is still isolated from them. It matches the comparison results in Table 2, where the order Leptospirales keeps a comparatively far distance from the other three orders by the phenotype standard. It indicates that the order Leptospirales should be promoted to an independent class in the phylum Spirochaetes or be separated from the phylum Spirochaetes.

### Planctomycetes

23 examples in the phylum Planctomycetes are included in the dataset and 1 species of them are separated from the main phylum part, connected to the phylum Kiritimatiellaeota. The species (class Phycisphaerae) is separated from the other species (class Planctomycetia) in phylum Planctomycete in the class level. Whereas, the species in the class Phycisphaerae locate very near to the class Planctomycetia by the “f_content” and “f_organization” models with Jaccard distance model (f_con_ja & f_org_ja).

Originally, the phylum Planctomycetes only contains one class, namely the class Planctomycetia [17]. Due to the reason that the species now in the class Phycisphaerae reproduce by binary fission, while other species in Planctomycetes reproduce by budding, a new class Phycisphaerae was proposed and separated from the class Planctomycetia [21,22].

To decrease the influence by single example in class Phycisphaerae, we utilize another 8 species (listed in SF3) in class Phycisphaerae for investigation and the partial MST result by org_ja method is shown in Figure 9. It is obviously that all species in phylum Planctomycetes are clustered into one part, and clearly separated into two groups, each representing one class. This result perfectly matches the NCBI taxonomy database.

**Figure 9.**
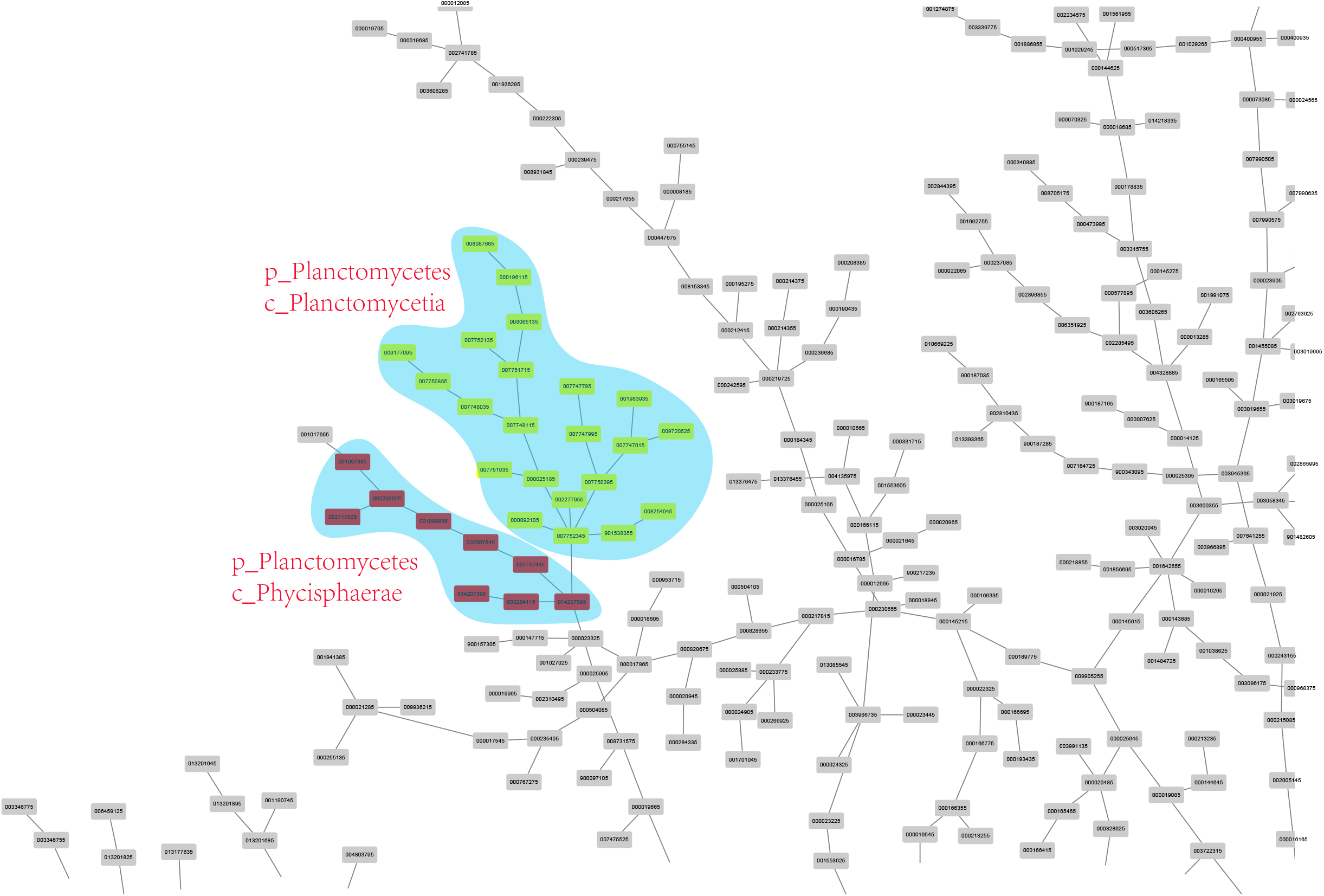
Updated MST result by adding another eight species in the class Phyciphaerae.

### Proteobacteria

1140 examples are included in the phylum and 1 species of them are far away separated from the main phylum part, being connected to the phylum Bacteroidetes, class Flavobacteriia. As for the other 1139 examples, they are either connected with each other or just located in a very near distance in the MST result. The 1 species (Glaciecola amylolytica) is divided from the other species of the phylum Proteobacteria in the species level. Since no other Glaciecola amylolytica examples is uploaded in the NCBI database. The situation remains as a problem to be solved in the future.

## Conclusion

In this paper, we proposed a framework to standardize the research on the taxonomy problem of bacteria based on protein domains. From the perspective of model evaluation, we find that the organization of the protein domains plays a more important role for bacterial taxonomy and phenotypes, compared with either the content of protein domain or the frequency of domains organization. Besides, the Jaccard distance can reflect a better relationship among all bacterial species than the Poisson distance and the Loss-corrected distance. From the perspective of taxonomy suggestions, we first proposed that the class Coriobacteriia should be removed from the phylum Actinobacteria and possibly into the phylum Firmicutes. Then, we suggest to move the order Acholeplasmatales of the phylum Tenericutes back to the phylum Firmicutes. Finally, we recommended that the order Leptospirales should be promoted to an independent class in the phylum Spirochaetes or separated from the phylum Spirochaetes.

## Acknowledgments

This work was supported by the National Natural Science Foundation of China [grant number 32070025, 31800136, 82041019]; the Research Project from State Key Laboratory of Pathogen and Biosecurity [grant number SKLPBS1807].

## Key Points

- A framework is proposed to normalize the process of bacterial taxonomy based on protein domain, which includes statistic model of domains, distance model, and construction and analysis models of taxonomy tree.
- By comparing the results produced by different methods, we find that the presence of the domain organization in a protein is most important in deciding the phenotypes of bacterial species.
- According to the best classification results by the framework, we proposed constructive suggestions on bacterial taxonomy.

## Figures and Tables

**Figure 1**. 3-step process in our proposed framework and candidates in each step.

**Figure 2**. An example of domains in one species and the corresponding records separately by four different statistic models.

**Figure 3**. A Venn diagram that shows concept of the corresponding three parts when comparing the domains of two species.

**Figure 4**. An example to construct the MST based on the fully connected network. (a) a fully connected network with distances marked. (b) A is first selected and then its nearest node B. (c) D has the nearest distance to A or B compared with C, and thus, involve D into the MST. (d) C is mostly near to D, and thus, connect C to D.

**Figure 5**. MST result by “content” model and Jaccard distance model. The MST result is manually preprocessed by grouping species in the phylum and class levels, and the phyla including more than 10 species are distinguished with different colors.

**Figure 6**. Twelve methods are compared by three different standards. Con, org, f_con, and f_org separately represent “content” model, “organization” model, “f_content” model, and “f_organization” model. Ja, po, and lo stand for Jaccard, Poisson, and Loss-corrected distance models, respectively.

**Figure 7**. Abstracted figure of MST generated by org_ja method. Six groups of species separated from their main phylum parts are connected to their neighboring phyla by the dashed line. Five of them are far away from their main phylum parts with yellow background and the numbers of species are also marked out.

**Figure 8**. Eight species in the order Leptospirales are added to generate the MST (left). Another nine species in Brachyspirales and one species in Brevinematales are involved to explore the relationship of the four orders in the phylum Spirochaetes (right).

**Figure 9**. Updated MST result by adding another eight species in the class Phyciphaerae.

## References

1. House C H, Fitz-Gibbon S T. Using homolog groups to create a whole-genomic tree of free-living organisms: an update[J]. Journal of molecular evolution, 2002, 54(4): 539–547.

2. Bansal A K, Meyer T E. Evolutionary analysis by whole-genome comparisons[J]. Journal of bacteriology, 2002, 184(8): 2260–2272.

3. Brown J R, Douady C J, Italia M J, et al. Universal trees based on large combined protein sequence data sets[J]. Nature genetics, 2001, 28(3): 281–285.

4. Baldauf S L, Roger A J, Wenk-Siefert I, et al. A kingdom-level phylogeny of eukaryotes based on combined protein data[J]. Science, 2000, 290(5493): 972–977.

5. Yang S, Doolittle R F, Bourne P E. Phylogeny determined by protein domain content[J]. Proceedings of the National Academy of Sciences, 2005, 102(2): 373–378.

6. Sarkar I, Gtari M, Tisa L S, et al. A novel phylogenetic tree based on the presence of protein domains in selected actinobacteria[J]. Antonie van Leeuwenhoek, 2019, 112(1): 101–107.

7. Fukami-Kobayashi K, Minezaki Y, Tateno Y, et al. A tree of life based on protein domain organizations[J]. Molecular biology and evolution, 2007, 24(5): 1181–1189.

8. Lytras S, Hughes J, Xia W, et al. Exploring the natural origins of SARS-CoV-2[J]. bioRxiv, 2021.

9. Wang Y, Zeng J, Zhang C, et al. New framework for recombination and adaptive evolution analysis with application to the novel coronavirus SARS-CoV-2[J]. Briefings in bioinformatics, 2021.

10. Finn R D, Bateman A, Clements J, et al. Pfam: the protein families database[J]. Nucleic acids research, 2014, 42(D1): D222–D230.

11. Letunic I, Bork P. Interactive Tree Of Life (iTOL) v5: an online tool for phylogenetic tree display and annotation[J]. Nucleic acids research, 2021, 49(W1): W293–W296.

12. Shannon P, Markiel A, Ozier O, et al. Cytoscape: a software environment for integrated models of biomolecular interaction networks[J]. Genome research, 2003, 13(11): 2498–2504.

13. Saitou N, Nei M. The neighbor-joining method: a new method for reconstructing phylogenetic trees[J]. Molecular biology and evolution, 1987, 4(4): 406–425.

14. Gao B, Gupta R S. Phylogenetic framework and molecular signatures for the main clades of the phylum Actinobacteria[J]. Microbiology and molecular biology reviews, 2012, 76(1): 66–112.

15. Gao B, Paramanathan R, Gupta R S. Signature proteins that are distinctive characteristics of Actinobacteria and their subgroups[J]. Antonie Van Leeuwenhoek, 2006, 90(1): 69–91.

16. Kunisawa T. Gene arrangements characteristic of the phylum Actinobacteria[J]. Antonie Van Leeuwenhoek, 2007, 92(3): 359–365.

17. Ludwig W, Euzéby J, Whitman W B. Road map of the phyla Bacteroidetes, Spirochaetes, Tenericutes (Mollicutes), Acidobacteria, Fibrobacteres, Fusobacteria, Dictyoglomi, Gemmatimonadetes, Lentisphaerae, Verrucomicrobia, Chlamydiae, and Planctomycetes[M] Bergey’s Manual® of Systematic Bacteriology. Springer, New York, NY, 2010: 1–19.

18. Brown D R, Bradbury J M, Johansson K E. Acholeplasmatales[J]. Bergey’s Manual of Systematics of Archaea and Bacteria, 2015: 1–2.

19. Karnachuk O V, Lukina A P, Kadnikov V V, et al. Targeted isolation based on metagenome - assembled genomes reveals a phylogenetically distinct group of thermophilic spirochetes from deep biosphere[J]. Environmental Microbiology, 2020.

20. Nakai R, Nishijima M, Tazato N, et al. Oligoflexus tunisiensis gen. nov., sp. nov., a Gram-negative, aerobic, filamentous bacterium of a novel proteobacterial lineage, and description of Oligoflexaceae fam. nov., Oligoflexales ord. nov. and Oligoflexia classis nov[J]. International journal of systematic and evolutionary microbiology, 2014, 64(Pt 10): 3353.

21. Fukunaga Y, Kurahashi M, Sakiyama Y, et al. Phycisphaera mikurensis gen. nov., sp. nov., isolated from a marine alga, and proposal of Phycisphaeraceae fam. nov., Phycisphaerales ord. nov. and Phycisphaerae classis nov. in the phylum Planctomycetes[J]. The Journal of general and applied microbiology, 2009, 55(4): 267–275.

22. Spring S, Bunk B, Spröer C, et al. Genome biology of a novel lineage of planctomycetes widespread in anoxic aquatic environments[J]. Environmental microbiology, 2018, 20(7): 2438–2455.

