## Supplementary figures and images for "Taxonomy Analysis in Bacterial Kingdom based on Protein Domain: A Comparison Study"

### Supplemental Figure 1

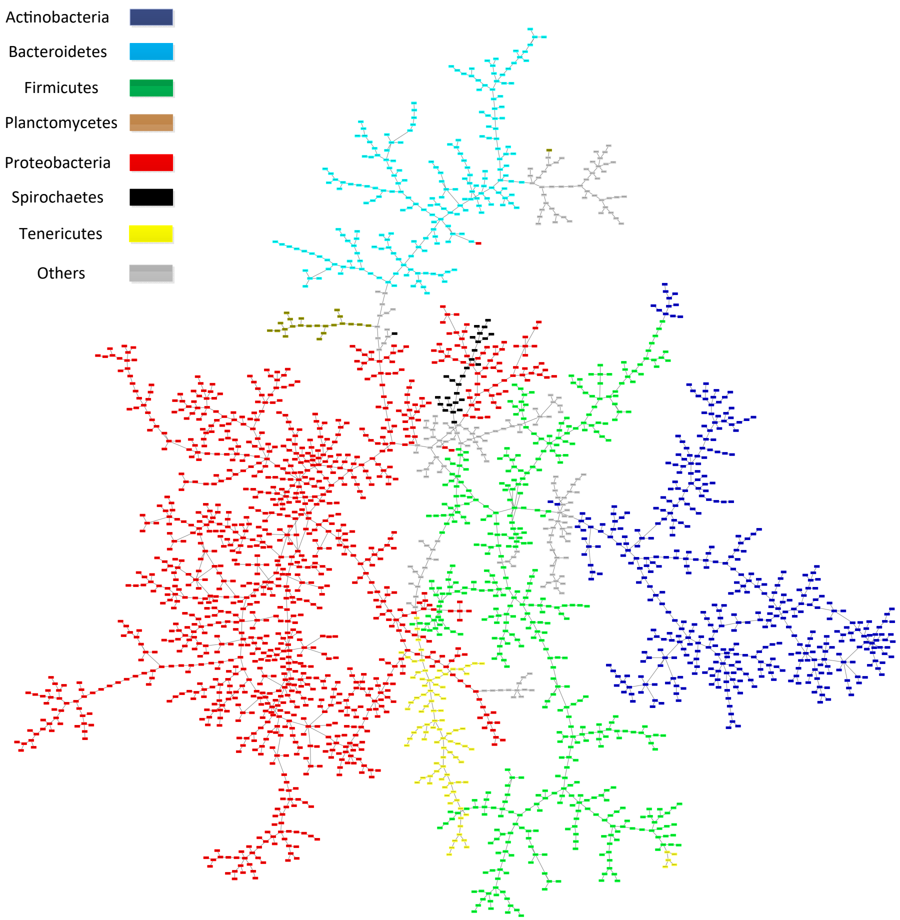


Supplementary Information Figure S1. MST result by "content" model and Jaccard distance.

### Supplemental Figure 2

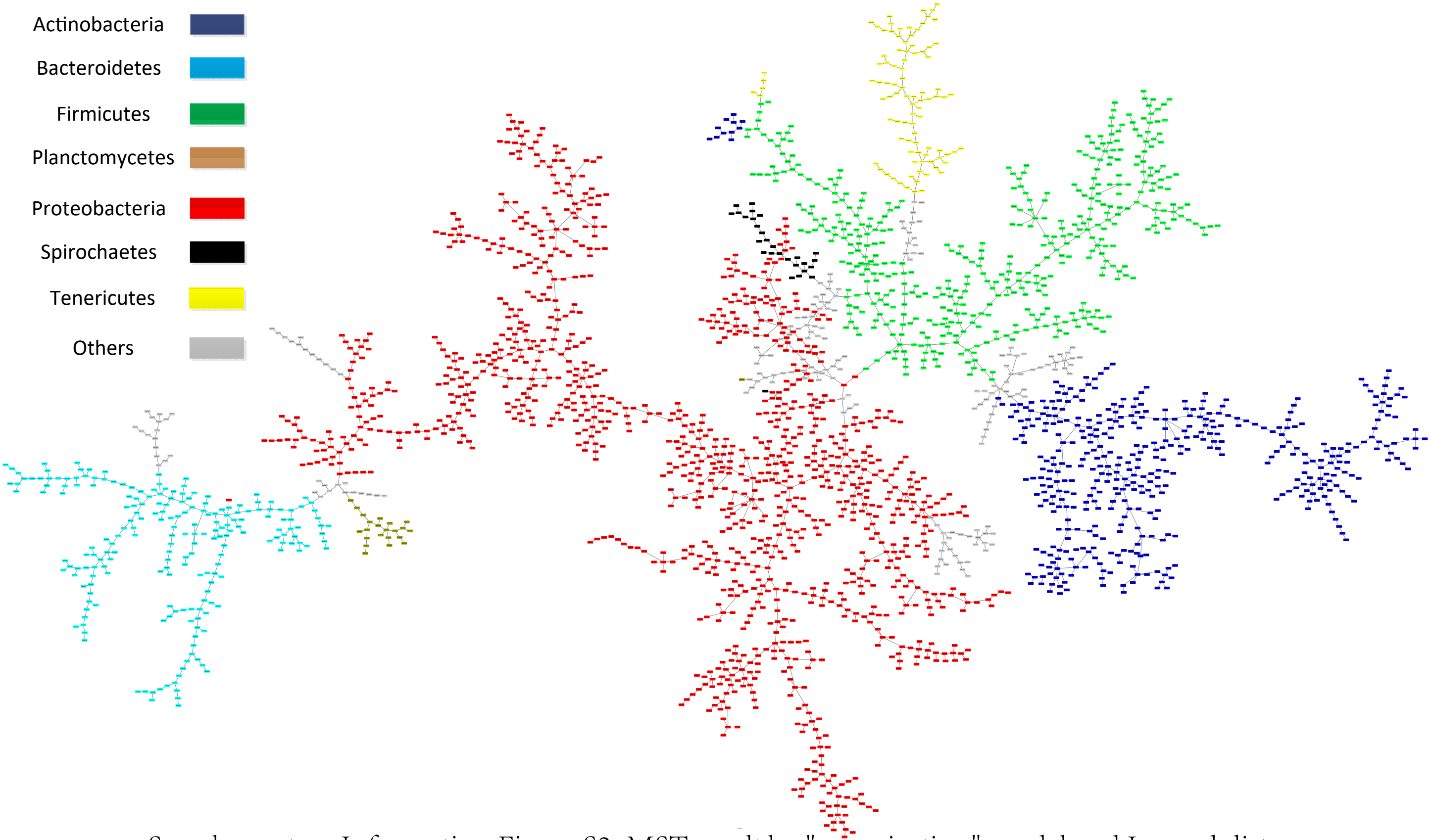


Supplementary Information Figure S2. MST result by "organization" model and Jaccard distance.

### Supplemental Figure 3

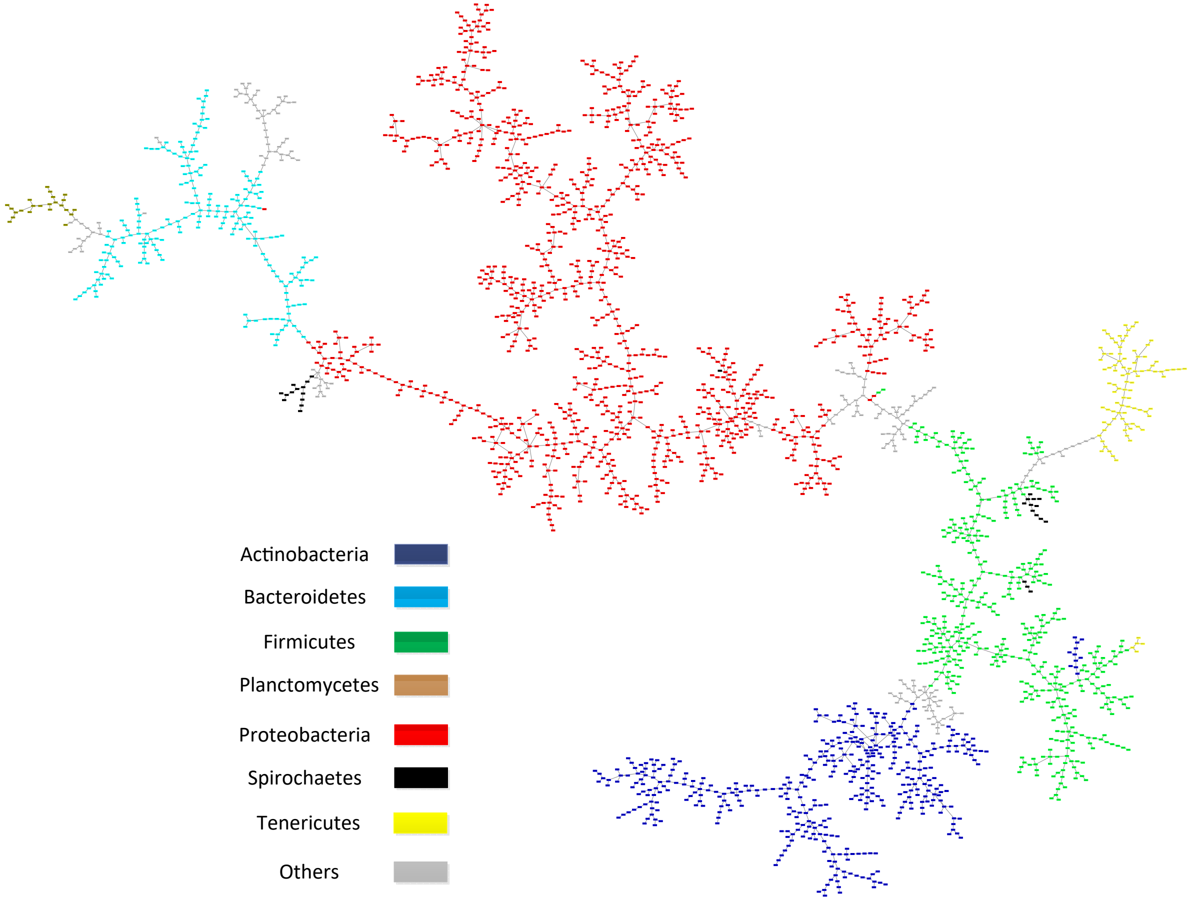


Supplementary Information Figure S3. MST result by "f_content" model and Jaccard distance.

### Supplemental Figure 4

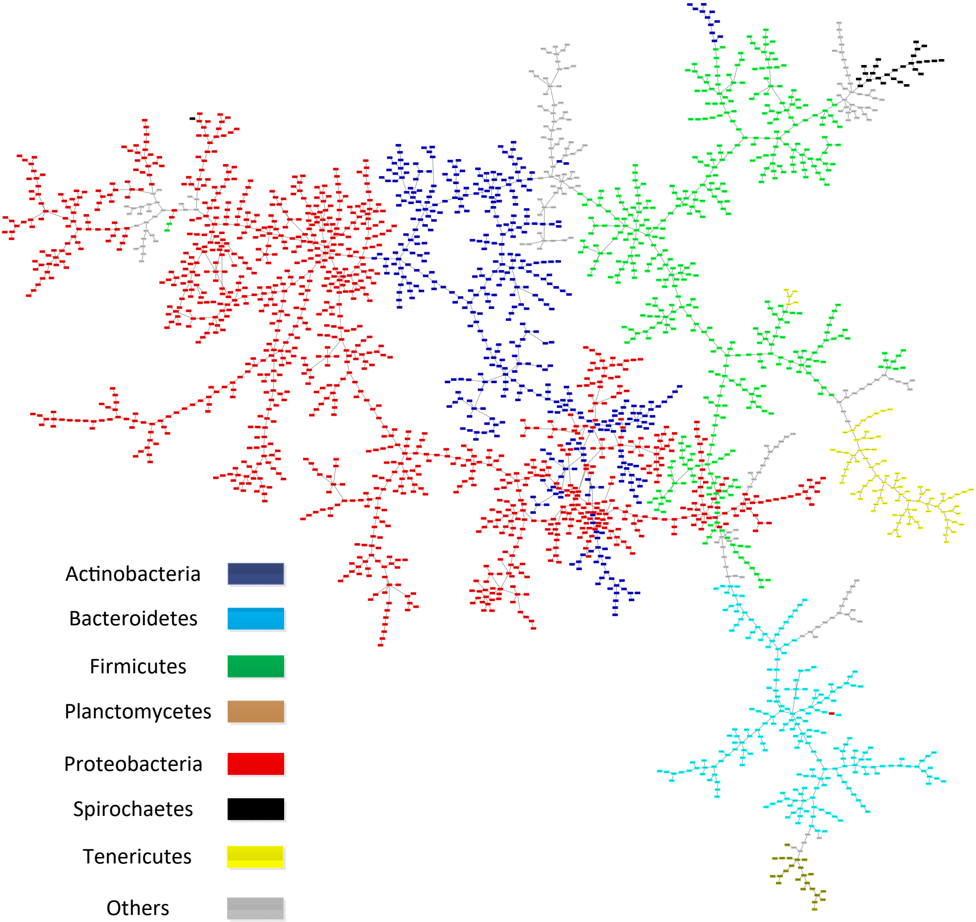

Supplementary Information Figure S4. MST result by "f_organization" model and Jaccard distance.

### Supplemental Figure 5

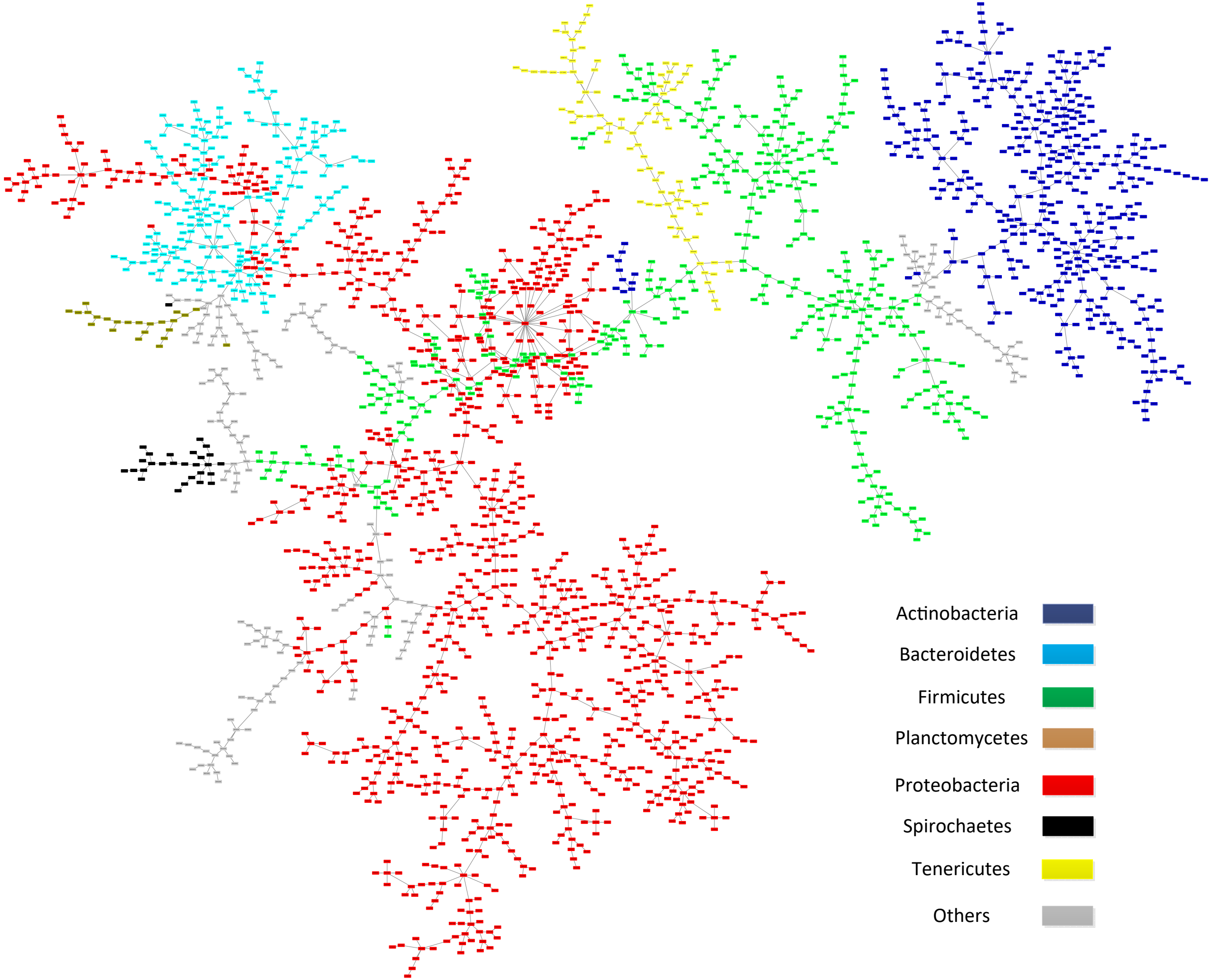


Supplementary Information Figure S5. MST result by "content" model and Poisson distance.

### Supplemental Figure 6

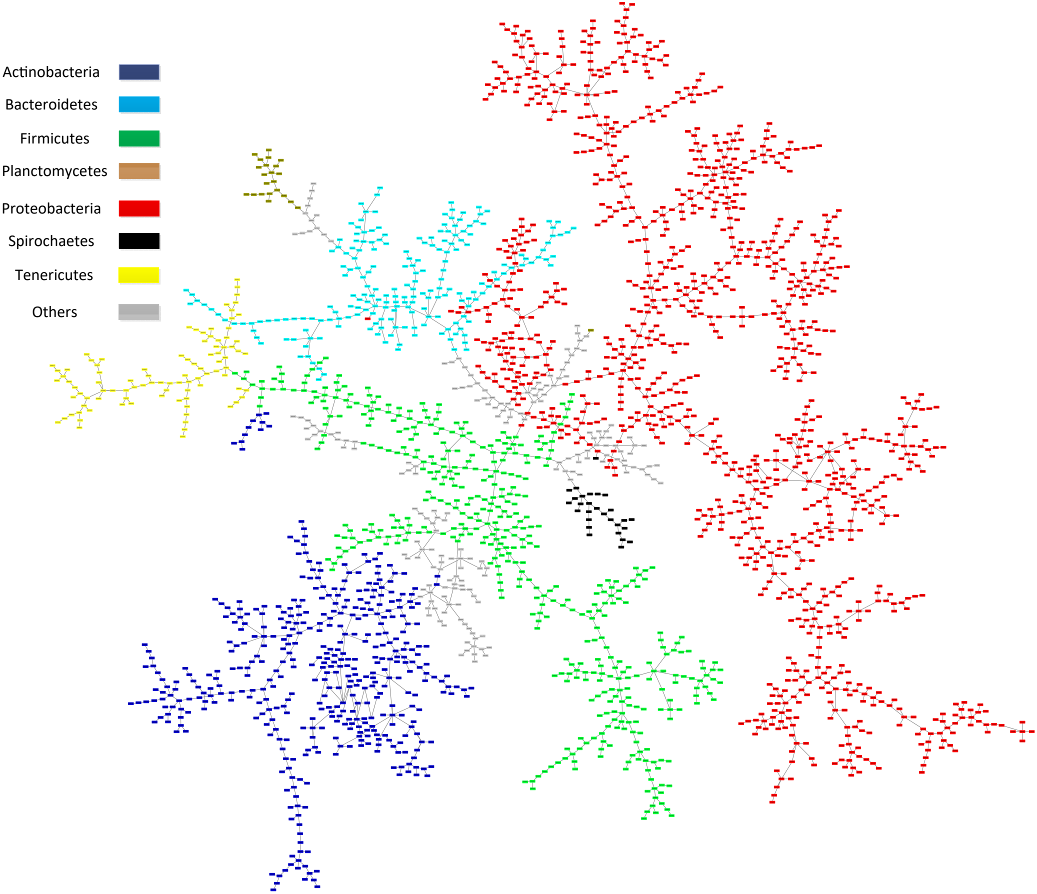


Supplementary Information Figure S6. MST result by "organization" model and Poisson distance.

### Supplemental Figure 7

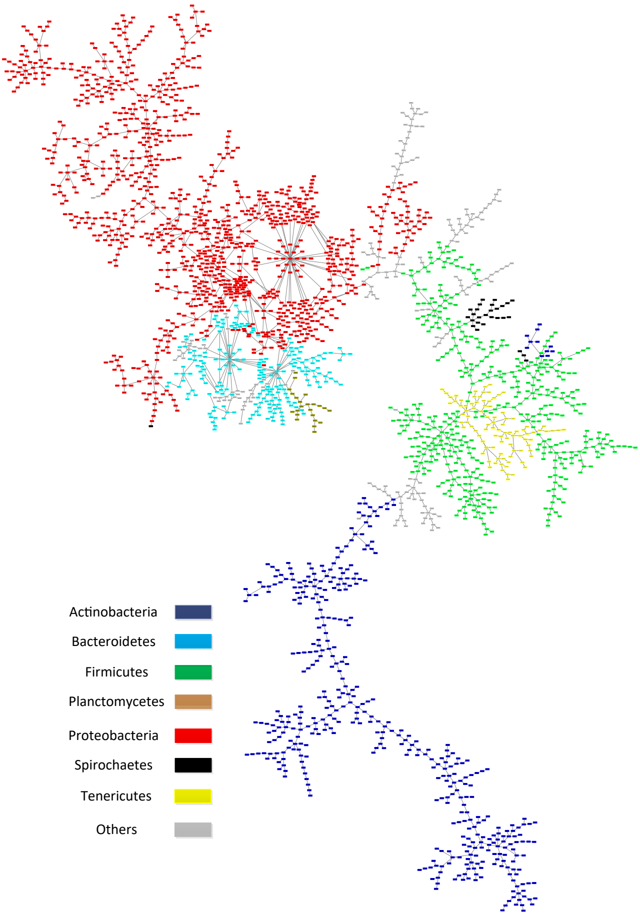


Supplementary Information Figure S7. MST result by "f_content" model and Poisson distance.

### Supplemental Figure 8

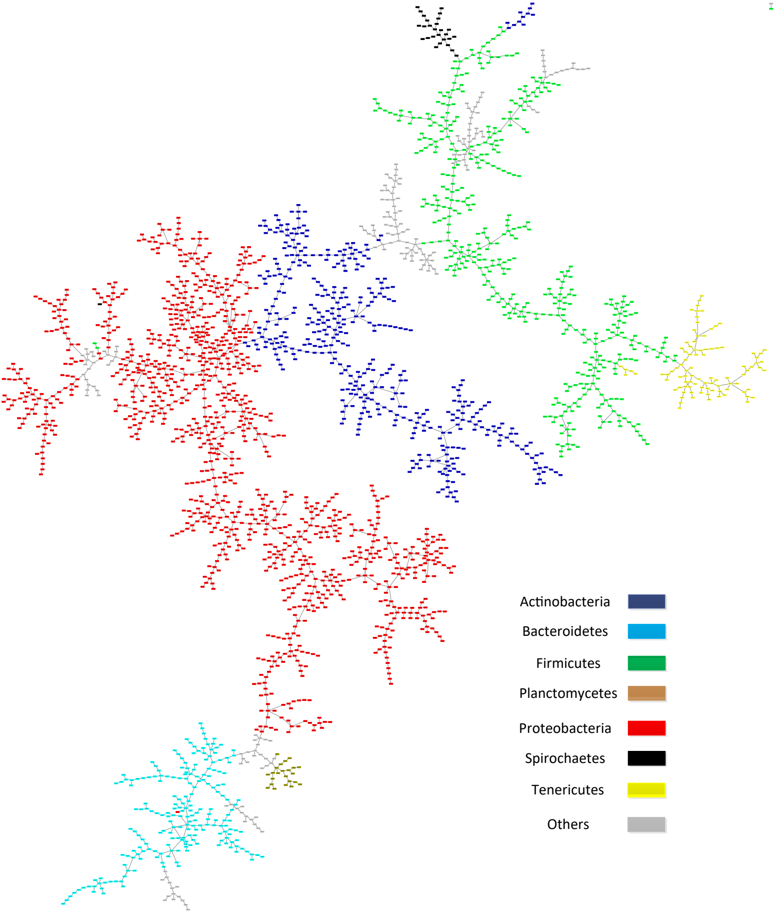


Supplementary Information Figure S8. MST result by "f_organization" model and Poisson distance.

### Supplemental Figure 9

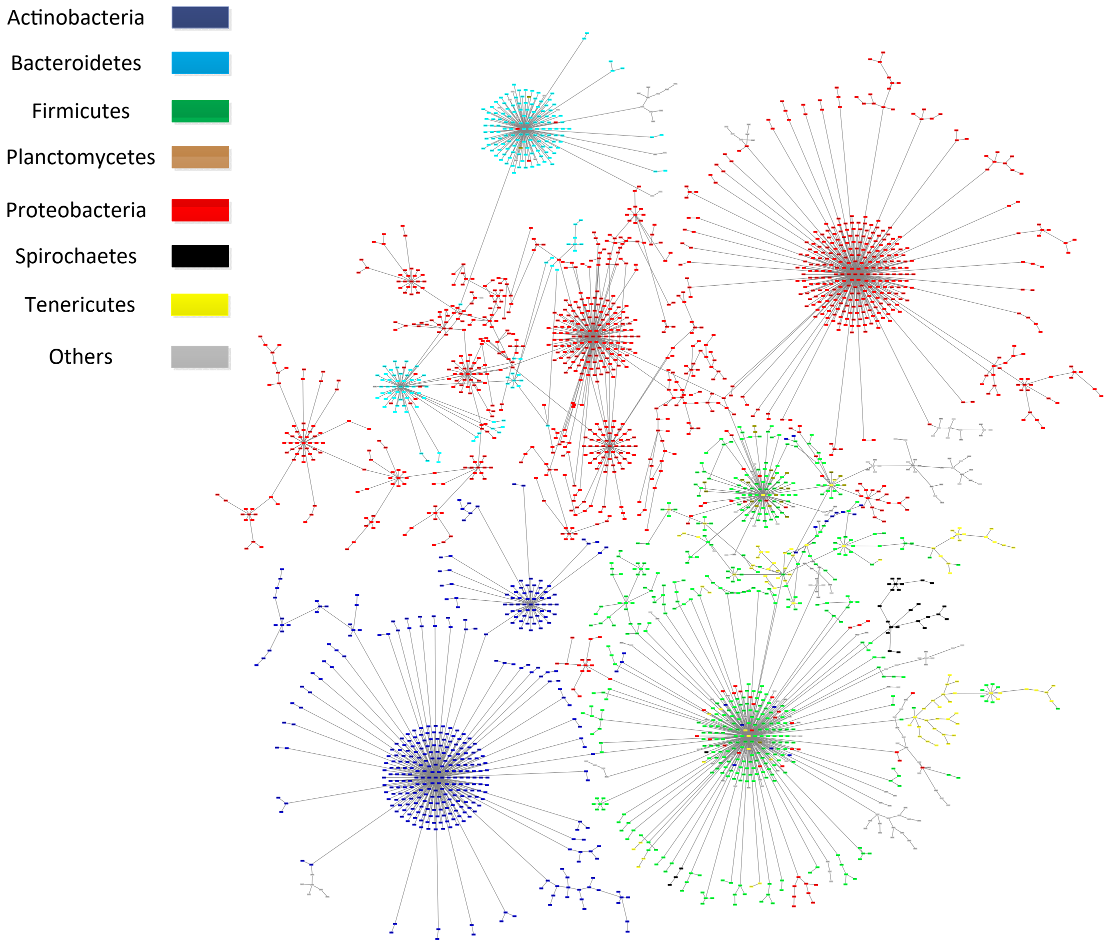


Supplementary Information Figure S9. MST result by "content" model and Loss-corrected distance.

### Supplemental Figure 11

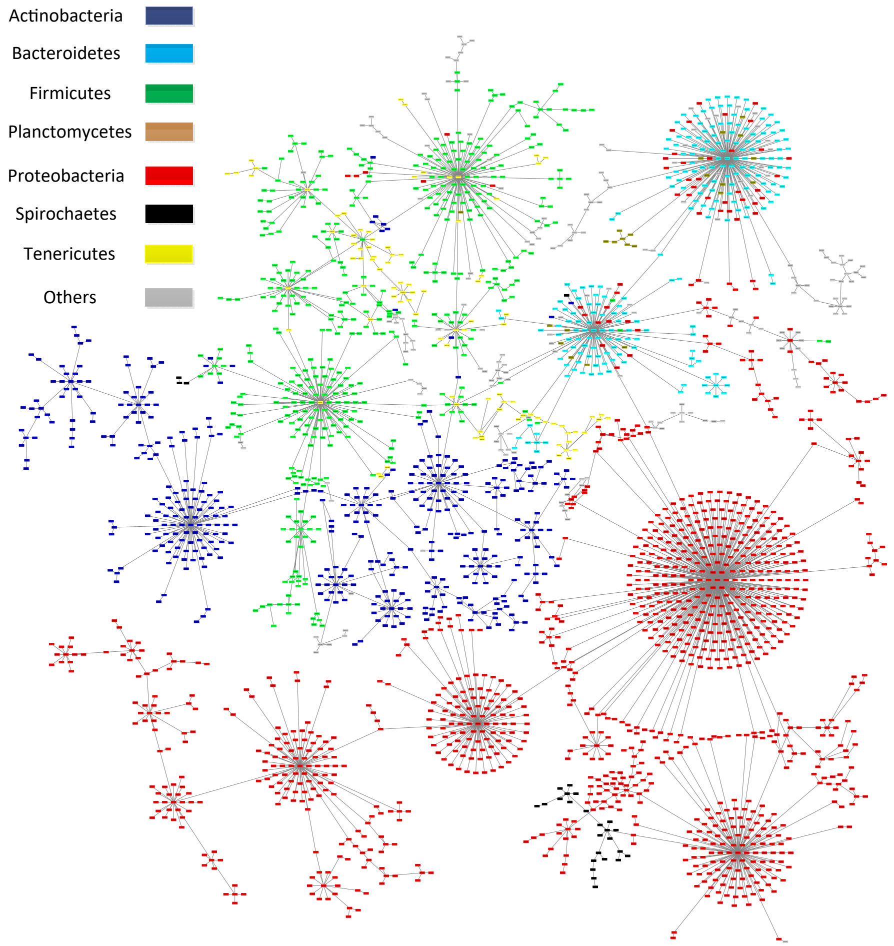


Supplementary Information Figure S11. MST result by "f_content" model and Loss-corrected distance.
