## Supplemental Figure 10 for "Taxonomy Analysis in Bacterial Kingdom based on Protein Domain: A Comparison Study"

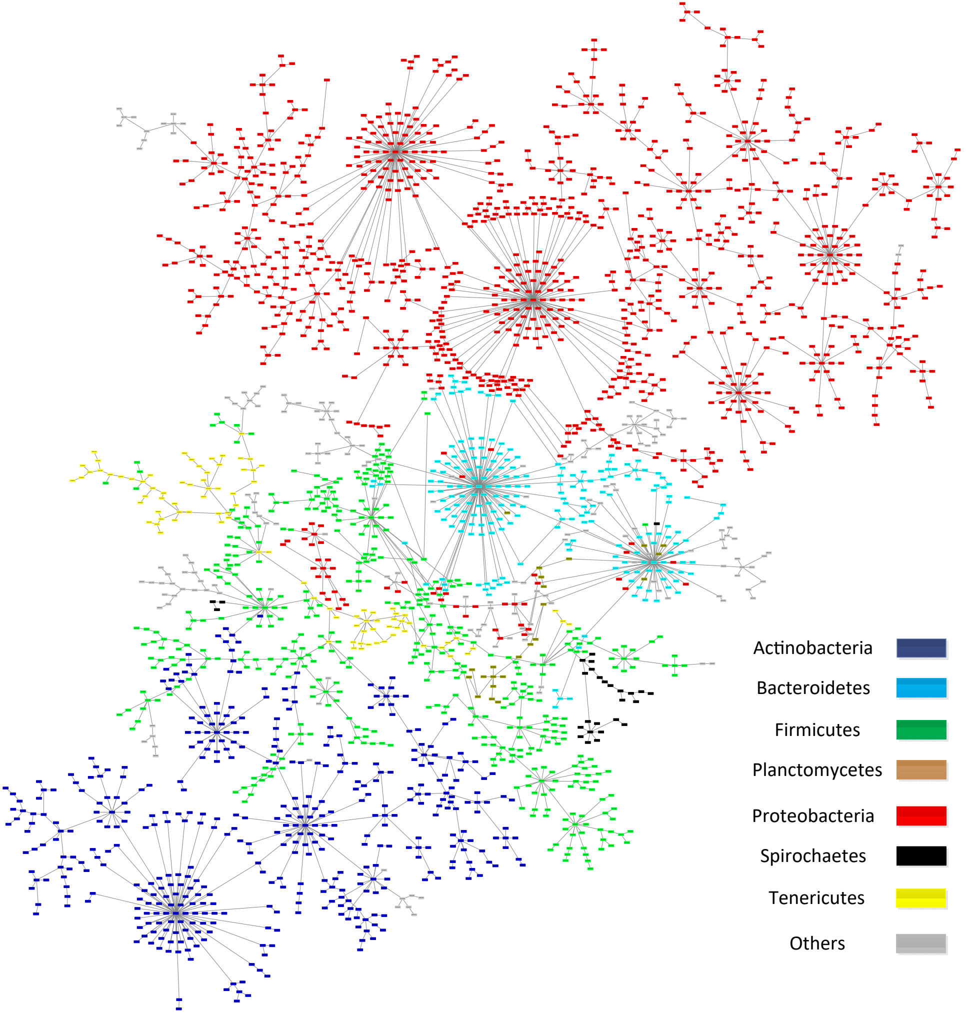


Supplementary Information Figure S10. MST result by "organization" model and Loss-corrected distance.
