## Supplemental Figure 12 for "Taxonomy Analysis in Bacterial Kingdom based on Protein Domain: A Comparison Study"

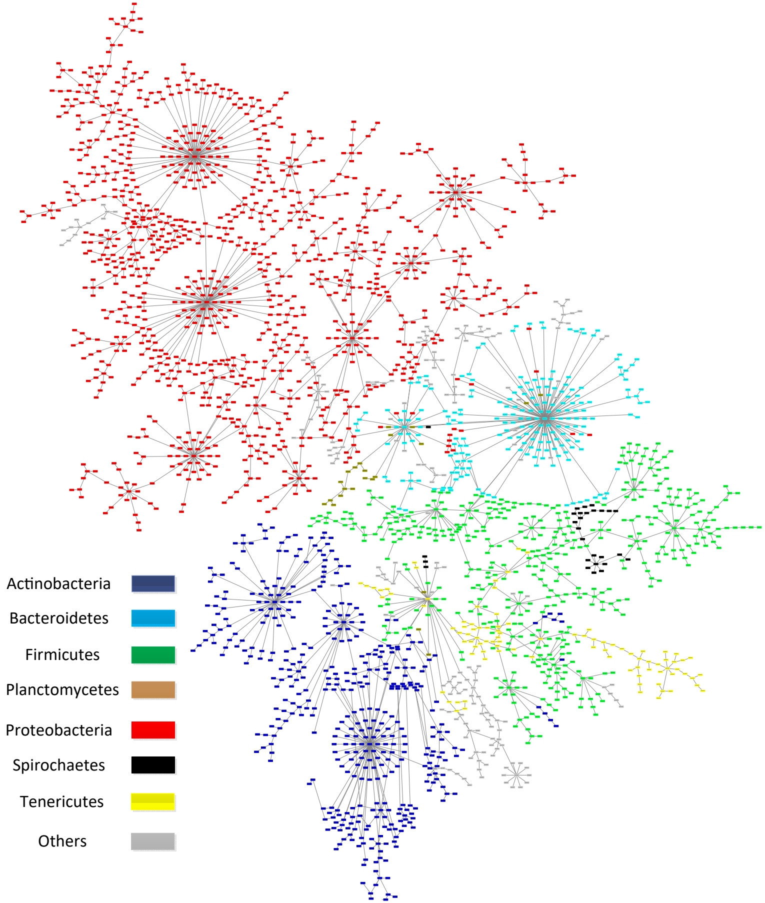


Supplementary Information Figure S12. MST result by "f_organization" model and Loss-corrected distance.
