## Supplemental Table 1 for "Taxonomy Analysis in Bacterial Kingdom based on Protein Domain: A Comparison Study"

Supplementary Information Table S1: Phyla classification results by Jaccard and Poisson distance models.

| | Phylum | total | con_ja | org_ja | f_con_ja | f_org_ja | con_po | org_po | f_con_po | f_org_po | common |
|---|---|---|---|---|---|---|---|---|---|---|---|
| 1 | Acidobacteria | 9 | 1 | | 1 | | 1 | | 1 | 1 | |
| 2 | Actinobacteria | 432 | 2, 11 | 2, 11 | 11 | 2, 11 | 2, 11 | 2, 11 | 11 | 2, 11 | 11 |
| 3 | Aquificae | 8 | | | | | | | | | |
| 4 | Archaea | 6 | | | | | | | | | |
| 5 | Armatimonadetes | 1 | | | | | | | | | |
| 6 | Bacteroidetes | 212 | | | | 1 | | | | | |
| 7 | Caldiserica | 1 | | | | | | | | | |
| 8 | Calditrichaeota | 1 | | | | | | | | | |
| 9 | Chlamydiae | 12 | | | | | | | | | |
| 10 | Chlorobi | 11 | | | | | | | | | |
| 11 | Chloroflexi | 13 | 3 | | 3 | 3 | 3 | 3 | 1, 3 | 3 | 3 |
| 12 | Coprothermobacterota | 1 | | | | | | | | | |
| 13 | Cyanobacteria | 28 | | | | | | | | | |
| 14 | Deferribacteres | 5 | | | | | | | | | |
| 15 | Deinococcus_Thermus | 24 | | | | | | | | | |
| 16 | Dictyoglomi | 2 | | | | | | | | | |
| 17 | Elusimicrobia | 2 | | | | | | | | | |
| 18 | Firmicutes | 458 | 2 | | 2, 3, 9 | 2, 9 | 2, 4, 150 | | 2, 4, 142, 150 | 2 | 2 |
| 19 | Fusobacteria | 15 | | | | | | | | | |
| 20 | Gemmatimonadetes | 3 | | | | | | | | | |
| 21 | Ignavibacteriae | 2 | | | | | | | | | |
| 22 | Kiritimatiellaeota | 1 | | | | | | | | | |
| 23 | Nitrospirae | 1 | | | | | | | | | |
| 24 | Planctomycetes | 23 | 1 | 1 | 1 | 1 | 1 | 1 | 1 | 1 | 1 |
| 25 | Proteobacteria | 1140 | 1, 2, 20, 74 | 1, 2, 12, 20, 27, 74 | 1, 2, 5, 53, 75 | 1, 2, 12, 20, 27, 74 | 1, 1, 1, 2, 3, 23,27 | 1, 2, 5, 12, 15, 27, 74 | 1, 2, 3, 23, 29 | 1, 2, 5, 27, 27, 74 | 1, 2 |
| 26 | Spirochaetes | 28 | 1 | 1 | 1, 3, 11 | 1 | 1 | 1 | 1, 3 | 1 | 1 |
| 27 | Synergistetes | 5 | | | | | | | | | |
| 28 | Tenericutes | 94 | 5 | 5 | 5 | 5 | | 5 | | 5 | 5 |
| 29 | Thermodesulfobacteria | 6 | | | | | | | | | |
| 30 | Thermotogae | 23 | | | | | | | | | |
| 31 | Verrucomicrobia | 7 | 2 | | 2, 2 | 3 | 2 | | 1, 2, 2 | 2 | 2 |
