## Supplemental Table 2 for "Taxonomy Analysis in Bacterial Kingdom based on Protein Domain: A Comparison Study"

Supplementary Information Table S2: Phyla classification results by Loss-corrected distance model.

| | Phylum | con_le | org_le | f_con_le | f_org_le |
|---|---|---|---|---|---|
| 1 | Acidobacteria | 7 | 4 | 8 | 4 |
| 2 | Actinobacteria | 13 | 5 | 11 | 5 |
| 3 | Aquificae | 4 | 2 | 4 | 3 |
| 4 | Archaea | 1 | 1 | 1 | 1 |
| 5 | Armatimonadetes | 1 | 1 | 1 | 1 |
| 6 | Bacteroidetes | 1 | 2 | 1 | 1 |
| 7 | Caldiserica | 1 | 1 | 1 | 1 |
| 8 | Calditrichaeota | 1 | 1 | 1 | 1 |
| 9 | Chlamydiae | 1 | 1 | 1 | 1 |
| 10 | Chlorobi | 2 | 2 | 3 | 2 |
| 11 | Chloroflexi | 10 | 7 | 9 | 6 |
| 12 | Coprothermobacterota | 1 | 1 | 1 | 1 |
| 13 | Cyanobacteria | 1 | 1 | 3 | 1 |
| 14 | Deferribacteres | 5 | 1 | 5 | 2 |
| 15 | Deinococcus_Thermus | 4 | 1 | 4 | 1 |
| 16 | Dictyoglomi | 1 | 1 | 1 | 1 |
| 17 | Elusimicrobia | 2 | 2 | 2 | 2 |
| 18 | Firmicutes | 307 | 53 | 291 | 37 |
| 19 | Fusobacteria | 9 | 2 | 6 | 2 |
| 20 | Gemmatimonadetes | 3 | 2 | 3 | 1 |
| 21 | Ignavibacteriae | 2 | 2 | 2 | 2 |
| 22 | Kiritimatiellaeota | 1 | 1 | 1 | 1 |
| 23 | Nitrospirae | 1 | 1 | 1 | 1 |
| 24 | Planctomycetes | 21 | 10 | 17 | 7 |
| 25 | Proteobacteria | 51 | 18 | 57 | 22 |
| 26 | Spirochaetes | 3 | 3 | 3 | 3 |
| 27 | Synergistetes | 4 | 1 | 4 | 1 |
| 28 | Tenericutes | 23 | 2 | 22 | 4 |
| 29 | Thermodesulfobacteria | 1 | 1 | 1 | 1 |
| 30 | Thermotogae | 5 | 1 | 5 | 3 |
| 31 | Verrucomicrobia | 5 | 5 | 6 | 5 |
